# Fly navigational responses to odor motion and gradient cues are tuned to plume statistics

**DOI:** 10.1101/2025.03.31.646361

**Authors:** Sam Brudner, Baohua Zhou, Viraaj Jayaram, Gustavo Madeira Santana, Damon A. Clark, Thierry Emonet

## Abstract

Odor cues guide animals to food and mates. Different environmental conditions can create differently patterned odor plumes, making navigation more challenging. Prior work has shown that animals turn upwind when they detect odor and cast crosswind when they lose it. Animals with bilateral olfactory sensors can also detect directional odor cues, such as odor gradient and odor motion. It remains unknown how animals use these two directional odor cues to guide crosswind navigation in odor plumes with distinct statistics. Here, we investigate this problem theoretically and experimentally. We show that these directional odor cues provide complementary information for navigation in different plume environments. We numerically analyzed real plumes to show that odor gradient cues are more informative about crosswind directions in relatively smooth odor plumes, while odor motion cues are more informative in turbulent or complex plumes. Neural networks trained to optimize crosswind turning converge to distinctive network structures that are tuned to odor gradient cues in smooth plumes and to odor motion cues in complex plumes. These trained networks improve the performance of artificial agents navigating plume environments that match the training environment. By recording *Drosophila* fruit flies as they navigated different odor plume environments, we verified that flies show the same correspondence between informative cues and plume types. Fly turning in the crosswind direction is correlated with odor gradients in smooth plumes and with odor motion in complex plumes. Overall, these results demonstrate that these directional odor cues are complementary across environments, and that animals exploit this relationship.

**Significance:** Many animals use smell to find food and mates, often navigating complex odor plumes shaped by environmental conditions. While upwind movement upon odor detection is well established, less is known about how animals steer crosswind to stay in the plume. We show that directional odor cues—gradients and motion—guide crosswind navigation differently depending on plume structure. Gradients carry more information in smooth plumes, while motion dominates in turbulent ones. Neural network trained to optimize crosswind navigation reflect this distinction, developing gradient sensitivity in smooth environments and motion sensitivity in complex ones. Experimentally, fruit flies adjust their turning behavior to prioritize the most informative cue in each context. These findings likely generalize to other animals navigating similarly structured odor plumes.

## Introduction

Many animals use their sense of smell to locate food and find mates [1-16]. To succeed in these tasks, they detect odor and wind patterns that inform their navigation choices (**Fig. 1A**) [17-19]. Multiple physical processes generate these sensory patterns. Odor molecules disperse from a source via molecular diffusion and are carried by the wind as odor plumes. Wind strength, turbulence, and other environmental factors affect the structure of these plumes and, consequently, the sensory patterns available to navigating animals [20, 21]. In steady air flows and near surfaces, odors form *smooth plumes*. In smooth plumes, molecular diffusion dominates and odor concentrations gradually decrease downwind from the source and decrease outward away from the plume’s centerline (**Fig. 1B)**[20, 22]. Under turbulent conditions, random air movements break up the odor fields into *complex plumes* composed of discrete filaments interspersed with odorless regions (**Fig 1C**) [3, 4, 8, 23-26]. In such complex plumes, the random movement of discrete odor filaments disperses odors from the centerline outward [27]. Across environments, the different physical processes that disperse odors laterally from the plume centerline lead to different plume statistics. There is evidence that statistical properties of visual scenes strongly influence how visual neural circuits process input signals [24, 28]. Different plume statistics could also convey different types of information for olfactory navigation, and navigators might rely on different cues in different environments. Here, we investigated this problem theoretically and empirically.

**Figure 1:**
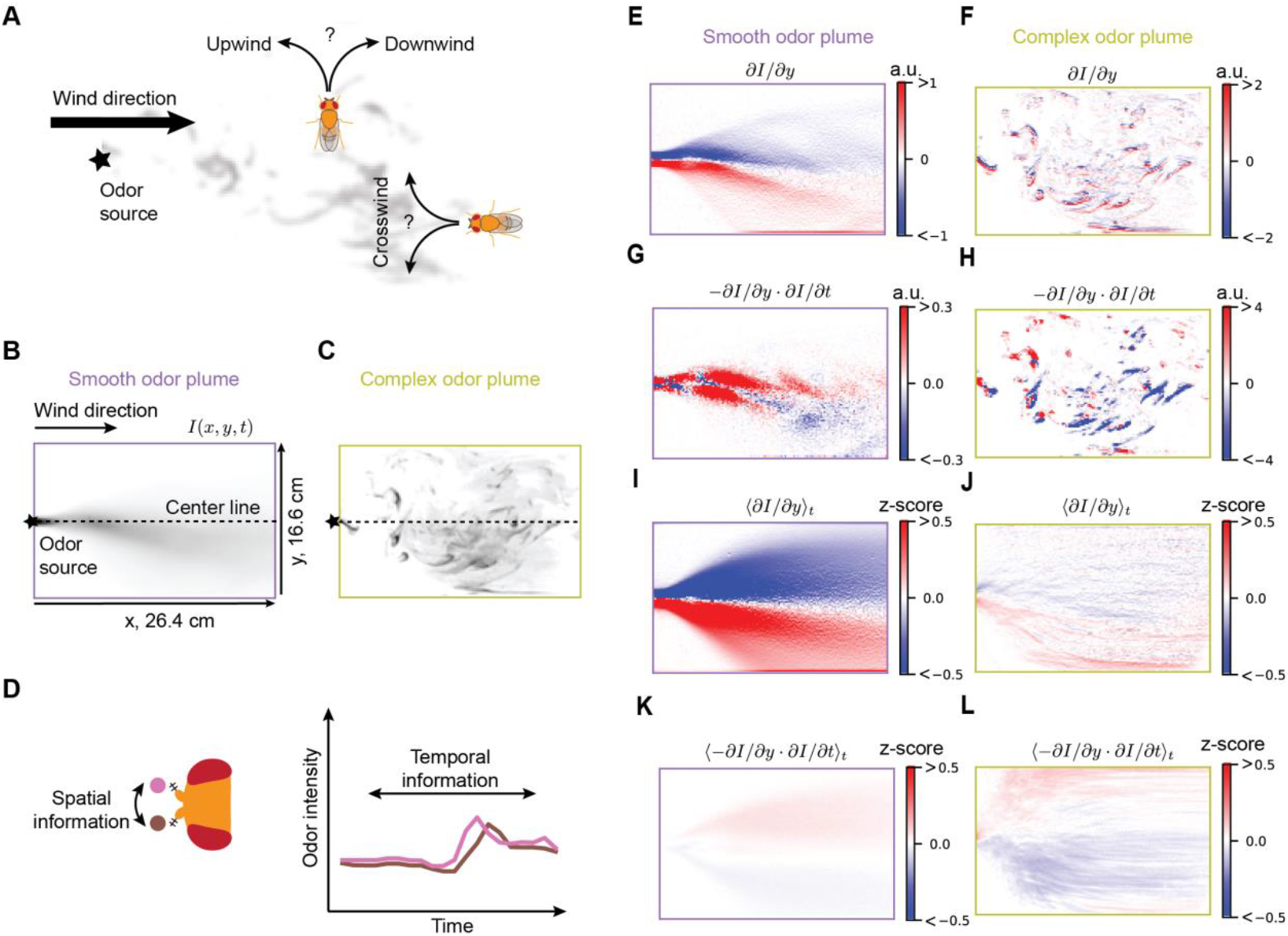
Directional information for navigation in different plume environments. (A) A fly needs to use various directional features to find the odor source (e.g., food). In a 2d environment, flies need to navigate both in the upwind/downwind and in the crosswind directions. Wind direction breaks symmetry in the upwind/downwind axis but does not distinguish crosswind directions. (B, C) Snapshots of two typical plume environments that a fly may encounter: a smooth odor plume and a complex odor plume. The odor intensity is represented by *I*(*x, y, t*). The upwind/downwind and crosswind axes are labeled *x* and *y*. (D) the fly’s two antennae can detect spatiotemporal information of the odor plume. (E, F) Gradient value in the *y* dimension in the two plume types. (G, H) Motion value in the *y* dimension in the two plume types (see Methods). (I, J) Time-averaged, z-scored gradient value in the *y* dimension. (K, L) Time-averaged, z-scored motion information in the *y* dimension (see Methods).

A basic strategy for navigating attractive odor plumes is to surge upwind when the odor is detected but cast crosswind when the signal is lost, allowing the animal to reconnect with the plume [3, 4, 22, 23, 29-40]. This surge-and-cast strategy has been extensively studied in theoretical models (Jayaram, Kadakia et al. 2022, Rigolli, Reddy et al. 2022, Jayaram, Sehdev et al. 2023, Verano, Panizon et al. 2023, Ouyang, True et al. 2024). A key feature of the strategy is that wind direction serves as the primary directional cue, while the odor signal itself only regulates when to surge upwind or cast crosswind. However, this approach has a significant limitation: although the wind direction serves as a key directional cue, it does not help an animal facing upwind distinguish between turns towards or away from the odor source (**Fig. 1A**). In other words, the wind breaks directional symmetry in the upwind-downwind axis but not in the crosswind direction, which is critical for efficiently locating the odor source.

Animals might improve on the surge-and-cast strategy by extracting directional information from the odor signal, which could complement information provided by the wind direction. In fact, many animals orient relative to directional olfactory cues that they calculate by comparing signals across separated bilateral sensors, like antennae in insects and nostrils in vertebrates. The spatial separation between sensors makes it possible to measure odor concentration at two locations and compare them. Stronger attractive stimulation of the left (right) side can drive left (right) turning in invertebrates [41-44] and vertebrates [14, 45], a response to the instantaneous spatial concentration difference, or *odor gradient* (**Fig. 1D**). In contrast, temporal comparisons between the right and left signals can provide information about the motion of the odor across the sensors, or *odor motion*. Experiments have shown that sequential activation of olfactory receptors in the left-to-right or right-to-left direction drive left (right) turning in invertebrates [7, 46] and vertebrates [9, 47], constituting responses to odor motion across the sensors (**Fig. 1D**).

These two directional odor cues are well-positioned to help navigators break symmetry perpendicular to the wind to distinguish between crosswind directions. In this way, they may help animals navigate toward plume centerlines (**Fig. 1A)** [7, 46]. Since odors can disperse from the centerline differently across environments, we wondered how plume structure would influence strategies to navigate crosswind using directional odor cues. Here we consider the roles of both odor gradient and odor motion signals in centerline-oriented navigation across plumes with different statistics. We hypothesized that odor gradients and odor motions would contribute differently to navigating environments with distinct statistics.

To test this idea, we analyzed the statistical properties of different odor plumes. We found that odor gradients and odor motions predict the direction to the plume centerline with different reliability in different plumes. We optimized shallow and interpretable neural network models to predict the centerline direction in each environment. These models showed differential sensitivity to odor gradient and motion cues that depended on their training environments. Using simulated agents, we demonstrated that using plume-appropriate directional odor cues improved navigation performance compared to using plume-inappropriate cues and compared to a strategy using wind as the only directional cue. Finally, we tracked *Drosophila* fruit flies navigating smooth and complex plumes and measured their crosswind navigation decisions. The fly behavioral responses to odor gradients and odor motion depended on the plume statistics and was consistent with a strategy that accounts for the differences in cue reliability across plumes. Overall, we show that optimized behavioral strategies exploit different directional odor cues in different environments and predict crosswind turning decisions by animals engaged in olfactory navigation.

## Results

### Directional odor features have plume-specific, antisymmetric organization around the centerline

We began by characterizing the spatiotemporal statistics of two recorded odor plumes generated under different conditions (**Fig. 1B, C**; see **Methods**) [20, 22, 46]. These plumes provide explicit information about how directional odor features – odor gradient and odor motion – are distributed across the centerlines. In the first recording, laminar airflow created a *smooth plume*, which has a smooth spatial distribution of odor intensity in time and space, *I*(*x, y, t*) [20, 22]. In the second recording, lateral perturbations of laminar flow create a sparser, *complex plume* with a more turbulent structure [46].

We hypothesized that differences in both odor gradients and odor motion distributions could be relevant for crosswind navigation. To assess differences in these distributions between plumes, we quantified the strength of the odor gradient and odor motion in the (cross-wind) *y* direction by calculating *∂I*/*∂y* and −*∂I*/*∂y* · *∂I*/*∂t*, respectively at each point (*x, y, t*) in each plume (snapshots in **Fig. 1E-H**; see Methods). In the remainder of the paper, we will refer to these as the gradient and motion features. These features were organized in qualitatively different ways in the smooth and complex plumes (**Fig. 1E-H**). In the smooth plume, the gradient feature had opposite signs on opposite sides of the centerline while in the complex plume, it did not exhibit this antisymmetric organization across the centerline. In contrast, the motion feature showed a different pattern of organization. In the smooth plume, there was no clear organization of the motion feature around the centerline, while in the complex plume the motion feature had opposite signs across the centerline. In summary, the gradient and motion features exhibited complementary distributions, with each organized anti-symmetrically around the centerline in one plume type but not the other.

Plume structure varies over time, so snapshots are not fully representative. Therefore, we calculated the temporal average of each feature at each location and divided these averages by the feature standard deviations to obtain z-scored values for each feature at each location (**Fig. 1I-L**). These z-scores can be interpreted as the signal-to-noise of the mean of each feature. The averages of both features exhibit antisymmetric organization around the plume centerline. However, the odor gradients have higher z-scores in the smooth plume than in the complex one, while the odor motion has higher z-scores in the complex plume than in the smooth one. These results suggest that, in smoother conditions, antisymmetric organization of the odor gradient is more reliable, while in more complex plumes, antisymmetric organization of the odor motion is more reliable.

### Plume-specific features predict centerline direction

If a feature is organized antisymmetrically across the plume centerline, it can provide information about the crosswind direction toward the centerline. We sought to quantify the predictive relationships between these features and the centerline direction for each plume. In each plume, we trained logistic regression models to predict the centerline direction using these odor features and subsequently scored their performance. The data were sampled randomly throughout each plume, assuming that the bilateral sensors faced upwind. They sampled two temporal odor signals, *R*(*t*) and *L*(*t*), at two nearby locations, 0.46mm apart, for a duration of 0.5s (**Fig. 2A**). We engineered three different features: the odor sum, the odor gradient, and the odor motion, defined as ⟨*L*(*t*) + *R*(*t*)⟩_*t*_, ⟨*L*(*t*) − *R*(*t*)⟩_*t*_, and ⟨*L*(*t* − Δ*t*)*R*(*t*) − *L*(*t*)*R*(*t* − Δ*t*)⟩_*t*_, respectively. The odor motion feature is the net spatiotemporal correlation in the odor intensity, with a delay of Δ*t* ≈ 0.017s, or one time step in our recordings (see Methods). Positive and negative net correlations indicate left-to-right and right-left odor motion. We trained our models to use these three features to best predict the probability that they are to the right or left of the centerline (**Fig. 2A**, see Methods).

**Figure 2:**
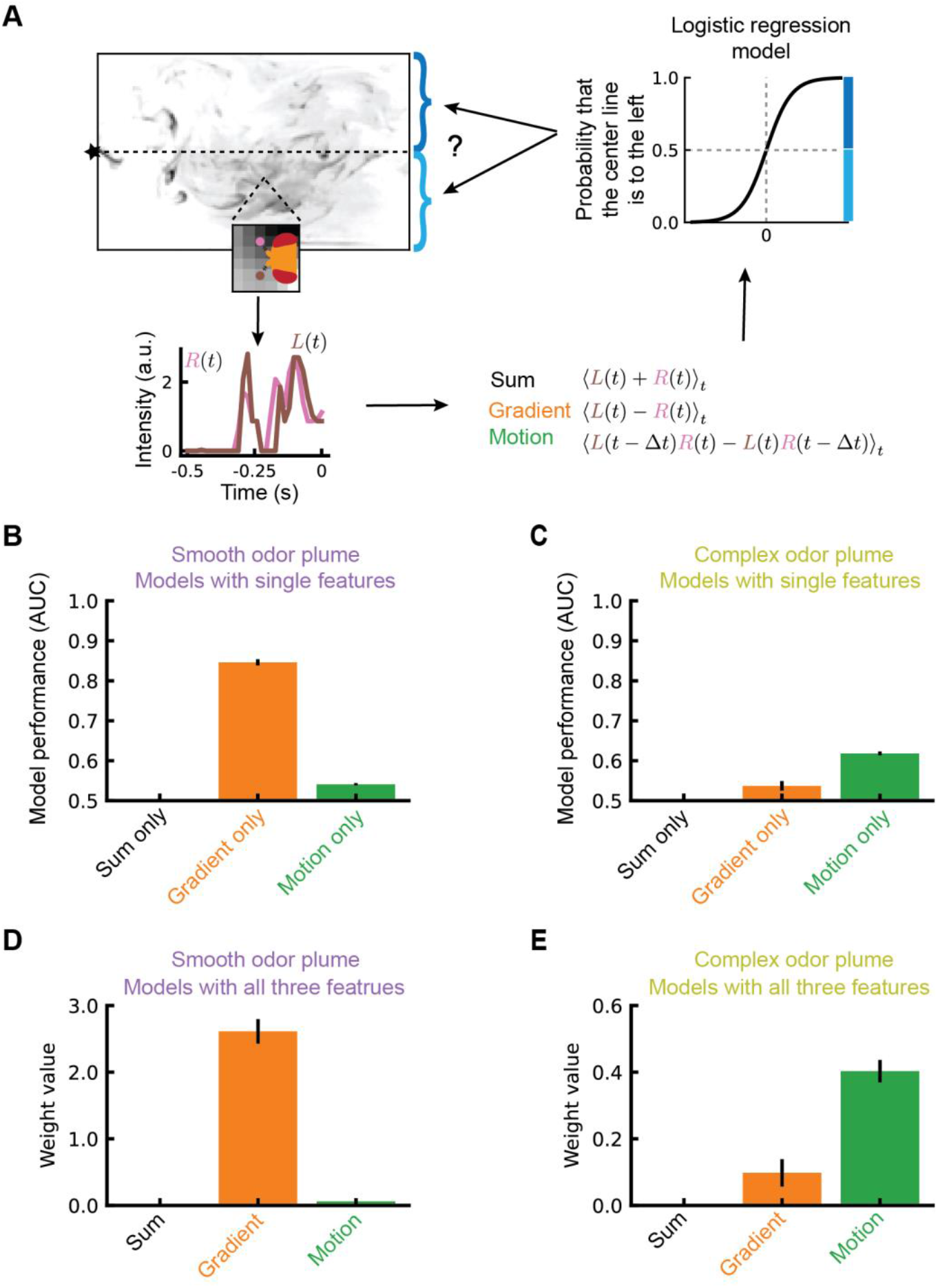
Gradient and motion information have different predictive powers in different plume types. (A) Three types of features are calculated from the two odor signals detected by the two antennae, *R*(*t*) and *L*(*t*): sum, gradient, and motion (see Methods). 500ms averages of feature traces are used to predict the direction to the centerline via logistic regression. (B, C) The three features are used individually to fit logistic regression models. Areas under the receiver operating curves (ROC AUC) are reported for the test samples. (D, E) The three features were used simultaneously to fit logistic regression models, and the trained weights are reported.

To assay the predictive power of each feature in each plume type, first we trained a separate model for each odor feature in each plume type. We assessed the performance of each of these models using receiver operating curves (ROCs), which reflect the tradeoff between detections and false positives for binary classifiers. ROCs can be summarized by the ROC area-under-the-curve (AUC) value, a statistic that is 0.5 for random classifier performance and approaches 1 for perfect classification. Models that used odor gradient to predict the smooth plume centerline direction had higher AUCs than models that used odor motion for the same task (**Fig. 2B, Fig. S2A**). This indicates that in the smooth plume, odor gradients were more predictive than odor motions of the centerline direction. In the complex plume, the opposite was true, so that odor motion was more predictive than odor gradients in determining the centerline direction (**Fig. 2C, Fig. S2B**). In both plume conditions, the bilateral sum ROC AUC values were 0.5, indicating that this control feature, which contains no spatiotemporal information, had no predictive power (**Fig. 2B, C**). In both plume types, odor motion features with one-frame delays had higher ROC AUC values than motion features using longer delays (**Fig. S2C**). Together, these quantifications support our previous qualitative description of the spatial distributions of odor gradient and odor motion features in the two plumes (**Fig. 1**). Specifically, in the smooth plume odor gradients but not motions reliably indicate the direction to the centerline. Conversely, in the complex plume odor motions but not gradients reliably indicate the centerline direction.

These single-feature models are conceptually simple, but they do not capture the possibility that correlations between gradient and motion features enable *both* to contribute to centerline predictions when these features are weighted and combined. To assess this possibility, we trained logistic models to use linear combinations of the z-scored values of all three features. In this analysis, a model’s fitted weight for a feature quantifies the influence of variation in that feature on classification. For example, a feature weight of zero magnitude indicates that the model ignores the effect of this feature. The model trained in the smooth plume learned a larger weight for the odor gradient feature than for the odor motion feature (**Fig. 2D, Fig. S2D**). Conversely, the model trained in the complex plume learned a larger weight for the odor motion feature than for the odor gradient feature (**Fig. 2E, Fig. S2D**). In both plumes, the weights for the odor sum feature were 0, as expected. These results indicate that, in different plume environments, odor gradient and motion features have different predictive powers about the centerline direction, even when considering correlations between the features.

### Optimized bilateral computation for centerline-finding recovers plume-specific odor features

Our analysis so far considered features that we engineered by computing averages and correlations between simulated antennal inputs. A real fly, however, does not have direct access to such features. To exploit the asymmetry of the gradient and motion features during olfactory navigation, a fly must process right and left antennal signals to extract relevant information. To investigate what such processing would look like, we built neural network models, which we trained to use bilateral timeseries to perform the same binary classification of the centerline direction as our logistic models above. After training, we assessed how solutions differed between the two plumes, focusing on the trained networks’ sensitivity to odor gradient and motion features.

First, we built a minimal network model (MNM) that has only two sets of trainable parameters, *f*_1_(*t*) and *f*_2_(*t*), the temporal filters acting on the odor signals at two simulated antennae (including an intercept parameter) (**Fig. 3A**). The MNM convolves these two temporal filters with the two odor signals, *R*(*t*) and *L*(*t*), sums these products, and passes the sum through a nonlinear activation function (ReLU). To incorporate left-right anti-symmetry in the model, we introduced an opponent structure to the model by subtracting a mirror-symmetric version of the same computation (**Fig. 3A**). Additionally, since real navigators make turns that alter the orientation of their sensors relative to their environment, the orientation of the artificial fly underwent a Wiener process but faced upwind at time 0, the most recent time in each training sample (See Methods; **Fig. S3A, B**), a technique that ensures that information in the distant past is not relevant to the inference[48, 49].

**Figure 3:**
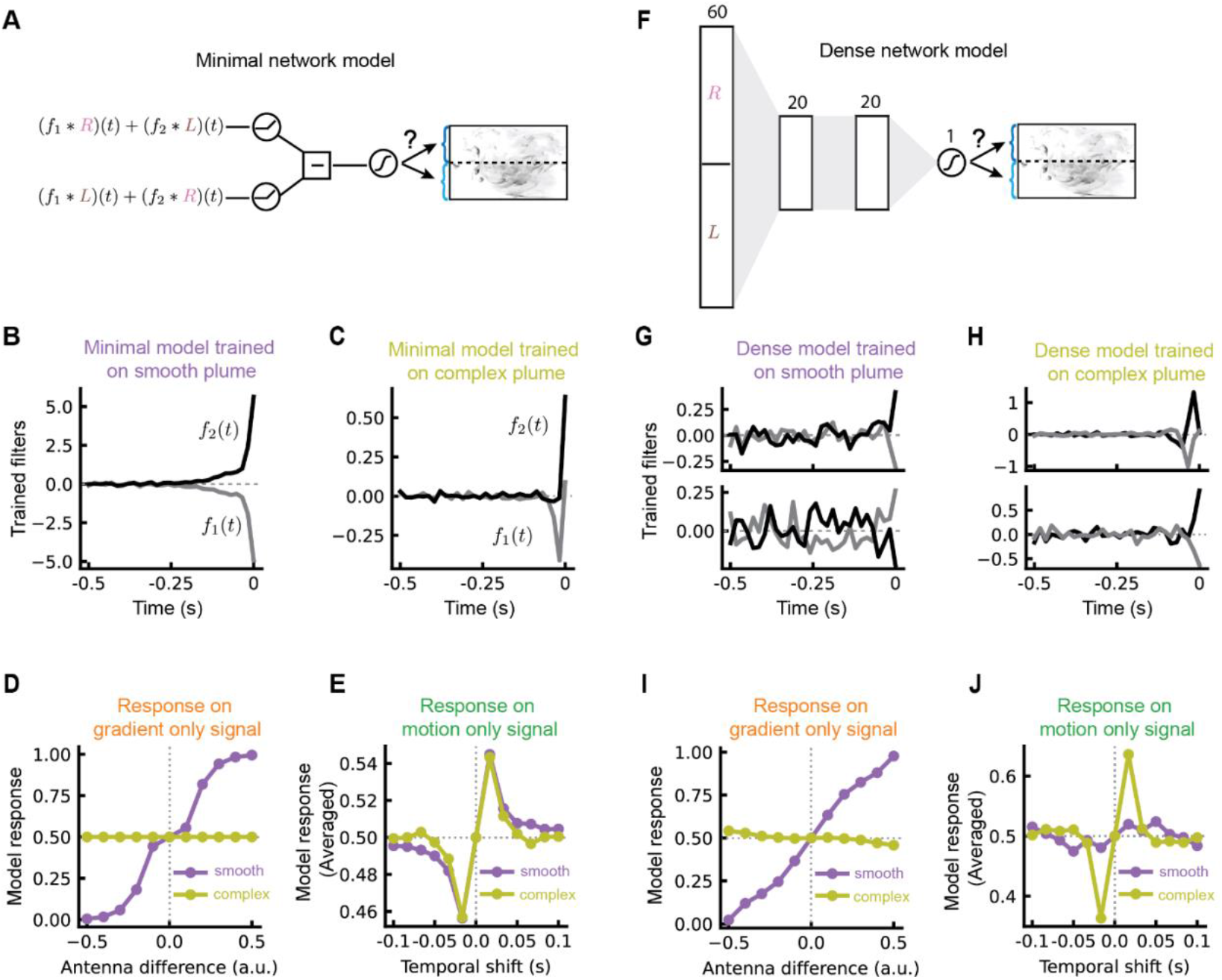
Neural network models trained in different plume types learn to sense different stimulus features. (A) A minimal network model (MNM) only has two trainable filters, represented by *f*_1_(*t*) and *f*_2_(*t*), respectively. The two inputs, *R*(*t*) and *L*(*t*), are generated when the fly head undergoes a rotational random walk (**Fig. S3A, B**). The output unit is a sigmoid function, which yields a probability that the centerline is to one side of the fly. (B) Trained filters of the MNM in the smooth odor plume, represented by the gray and black lines, respectively. (C) Trained filters of the MNM in the complex odor plume, represented by the gray and black lines, respectively. (D) Responses of the trained MNMs to gradient stimuli as a function of the antennal difference (see Methods). (E) Responses of the trained MNMs to motion stimuli as a function of the temporal shift between the two input signals (see Methods). (F) A dense network model (DNM) has two hidden layers, each with 20 units. The feedforward connections between the input layers (by concatenating the two inputs), the first hidden layer, and the second hidden layer are all-to-all. The output unit is a sigmoid function, which gives a probability that the centerline is to one side of the fly. (G) Examples of first layer filters trained in the smooth plume between the input layer and the first hidden layer (2 out of 20). The black lines correspond to the filters acting on *L*(*t*) while the grey lines correspond to the filters acting on *R*(*t*). (H) As in (G), but for filters trained in the complex plume. (I) As in (D), but for the DNMs. (J) As in (E), but for the DNMs.

We trained two MNMs to identify the centerline direction in each plume type (**Fig. S3C-E**). The resulting filters had very different structures in the two plumes. For models trained in the smooth plume, the filters acted like a gradient detector (**Fig. 3B**), computing the difference in intensity between the two antennae. For models trained in the complex plume, filters were more complex (**Fig. 3C**). The opposite signs of the filters suggest that they are also sensitive to gradients, but the delayed, negative lobe in *f*_1_(*t*) is reminiscent of a visual motion detection model proposed based on data in the rabbit retina [50], for which the delayed inhibition in *f*_1_(*t*) tends to veto signals when they arrive in the non-preferred order.

To better quantify the response properties of each model, we created synthetic signals containing exclusively gradient or motion content (**Fig. S3F, G**). For the gradient content, we varied the difference in intensity between the two antennal inputs (see Methods). For the motion content, we generated binary signals and enforced a correlation between the two antenna signals with different delays (see Methods) [51]. The model trained in the smooth plume responded more strongly to the gradient signals than the model trained in the complex plume (**Fig. 3D**). Interestingly, the models trained in the complex and smooth plumes both responded strongly to the motion signals with various delays (**Fig. 3E**). The gradient detector obtained from training in the smooth plume responds to the motion signals because of small temporal asymmetries between the two filters (**Fig. 3B**). Responses to the motion signals disappear if exactly symmetrical filters are used (**Fig. S3H**).

The minimal models are interpretable, but not very expressive. More expressive models could allow us to assay the behavior of algorithms—particularly their responses to gradients and motion—closer to the upper limit of task performance. We therefore built a general dense network model (DNM) with the same opponency structure (see Methods) to predict the centerline direction (**Fig. 3F, Fig. S3I-K**). This dense model employs 20 sets of temporal filters between the input layer and the first hidden layer (**Fig. 3G, H**), but only performs modestly better than the MNM (**Fig. S3K**). Interestingly, the results with these more expressive models are similar to the results with the MNM. The temporal filters trained in the smooth plume look more like gradient detectors, while those trained in the complex plume look more like motion detectors, although in some cases the distinction was less clear (bottom row in **Fig. 3G, H**). To evaluate the sensitivity of these models to odor gradients and odor motion, we tested the trained DNMs on the same synthetic data as before (**Fig. S3F, G**). When tested on the gradient signals, the model trained in the smooth plume was more sensitive to gradient signals compared to the model trained in the complex plume (**Fig. 3I**), similar to the results with MNMs. On the other hand, while the MNMs trained in the complex and smooth plumes responded to the motion signals with similar strength, the DNM trained in the complex plumes responded more strongly to the motion signals than the one trained in the smooth plumes (**Fig. 3J**). These results suggest that increased flexibility afforded by the dense architecture allowed the models to become even more specialized for gradient and motion features when trained in the smooth and complex plume respectively.

### Sensitivity to plume-appropriate directional features enhances agent navigation

Our previous results indicate that, in different plumes, different computations are useful for predicting the centerline direction. To support navigation, responses to these features must be integrated with broader strategies to reach the odor source. Critically, they must be combined with odor timing cues that indicate the distance, but not the direction, to the plume centerline [33, 52], and that regulate turning in the upwind or downwind direction [22, 33, 52]. We hypothesized that navigation strategies incorporating directional odor information would outperform strategies based only on odor timing and the wind direction. Specifically, we hypothesized that these improvements would depend on the same correspondence between specific directional odor features and the plume types that we found in our earlier analysis of centerline prediction. To test this idea, we built artificial agents that navigated the two plumes using different odor directional features while quantifying the agents’ ability to find the odor source.

Our simulated agents navigated odor plumes by alternating straight forward movement with instantaneous turns. Turns occurred stochastically at a constant rate (4/3 Hz). When turns occurred, their magnitude was normally distributed with a mean of 30° and standard deviation of 8°, following previous models of fly turning [33]. While agents had direct knowledge of the average wind flow direction, information about whether the plume centerline was on their left or right was not provided to them directly (**Fig. 4A**). Instead, agents used the wind direction and bilateral odor features to choose between clockwise and counterclockwise turning according to different strategies (**Fig. 4A-B, Fig. S4A**). The baseline model used only wind direction, while the probability of upwind or downwind turning depended dynamically on the sum of the left and right antennae signals, (**Fig. S4B-D** and Methods), based on a published model of odor-gated upwind turning in flies [32]. In the augmented strategies, agents also estimated the direction to plume centerline by passing their bilateral odor histories to the DNM trained on either the smooth plume or the complex plume (**Fig. 4B**). If the network provided only weak evidence about the centerline direction, these agents used the baseline strategy. If the network provided strong information about the centerline direction, the agents turned left or right according to the estimated direction to centerline (**Fig. 4B**).

**Figure 4:**
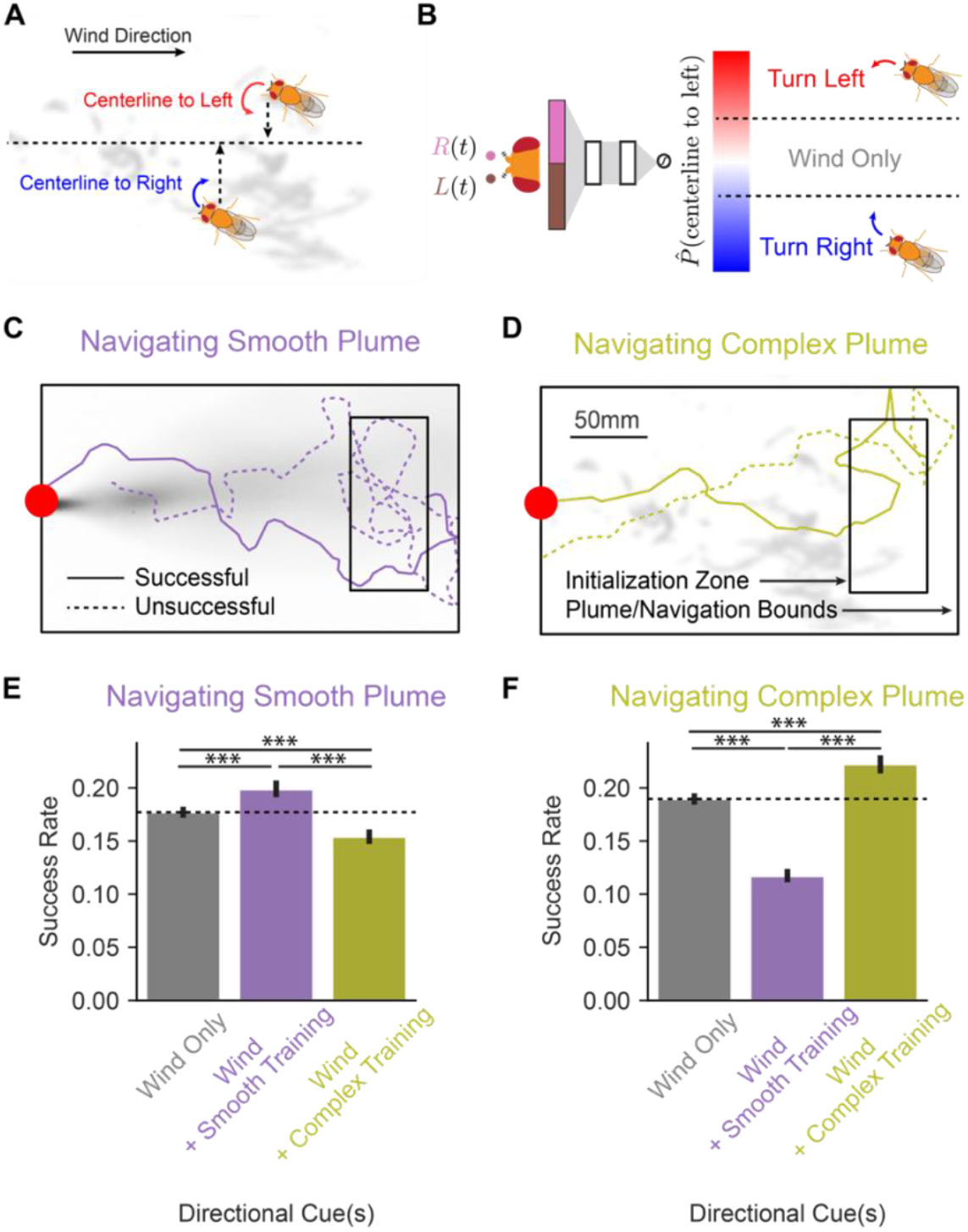
Sensitivity to plume-appropriate directional odor features improves wind-based olfactory navigation strategies. (A) Simulated agents navigated smooth or complex plumes. Navigation choices were made using the wind direction and estimates of the direction to the plume centerline. (B) Bilateral signals detected by the agent were passed through a network model (*left*, see **Fig. 3**) to obtain an estimate of the probability that the centerline was to the agent’s left or the agent’s right. If the network model only weakly predicted the centerline direction, agents turned upwind or downwind according to a previously published model that does not account for bilateral directional cues. If network prediction strength exceeded a threshold, agents turned towards the estimated direction to centerline. (C-D) Example successful and unsuccessful trajectories of agents navigating the smooth (C) or complex (D) plume, using the smooth-trained or complex-trained network model respectively. All agents were initialized in a box located in the crosswind plume center, with a 112mm width and extending 200-250mm downwind of the source (10mm radius, red dot). Successful agents reached a 10mm-radius goal at the plume’s source. Unsuccessful agents either reached a maximum of 90s search time without reaching the source (C) or traveled upwind beyond the goal without entering it (D). (E-F) For agents navigating the smooth plume (E), or complex plume (F), success rates for baseline navigators lacking bilateral features (*gray*), with a bilateral, centerline-predicting network trained in the smooth plume (*purple*), or with a bilateral, centerline-predicting network trained in the complex plume (*gold*). Error bars are 95% confidence intervals. Pairwise comparisons of success rate assessed by z-tests (n=10,000 trajectories per condition; *** indicates p < 0.001).

All agents were initialized randomly inside a box near the back of the plume (**Fig. 4C-D**). They were considered successful if they entered a 10mm radius circle at the odor source (**Fig. 4C-D**). If they failed to enter this region in 90 seconds, or if they walked upwind past the source, they were considered unsuccessful. In the smooth-plume, agents using the augmented strategy with the smooth plume-trained dense network outperformed agents using the baseline strategy (**Fig. 4E**). Similarly, in the complex plume, agents using the augmented strategy with the complex plume-trained dense network also outperformed agents using the baseline strategy (**Fig. 4F**). However, agents using the augmented strategy with plume-inappropriate computations—either smooth plume-trained networks during complex plume navigation or complex plume-trained networks during smooth plume navigation—performed even worse than agents using the baseline strategy. Across both plume types, bilateral computations predicting the centerline enabled more successful navigation than strategies based only on odor timing and the wind direction; however, using a plume-appropriate computation was critical to this improvement.

### Different directional odor cues predict fly crosswind turning in different plume types

Bilateral gradient and motion cues drive orienting responses in fruit flies [41-43, 46, 53] and other animals [7, 9, 13, 14, 47]. After observing that these cues have different utility across plume types, we wondered whether animal behavioral responses to these cues depend on their navigation environment. Specifically, we sought to quantify the relationship between these cues and crosswind turning in flies navigating smooth and complex odor plumes under otherwise identical conditions. To present the plumes precisely, we used structured optogenetic olfactory stimulation instead of real odors, which are difficult to control. We optically projected our model plumes onto a navigation arena while flowing laminar air parallel to the plume centerline (**Fig. 5A**)[46]. We introduced flies into the arena that express CsChrimson in their olfactory neurons (see Methods), thereby making the olfactory neurons light sensitive [32, 46].

**Figure 5:**
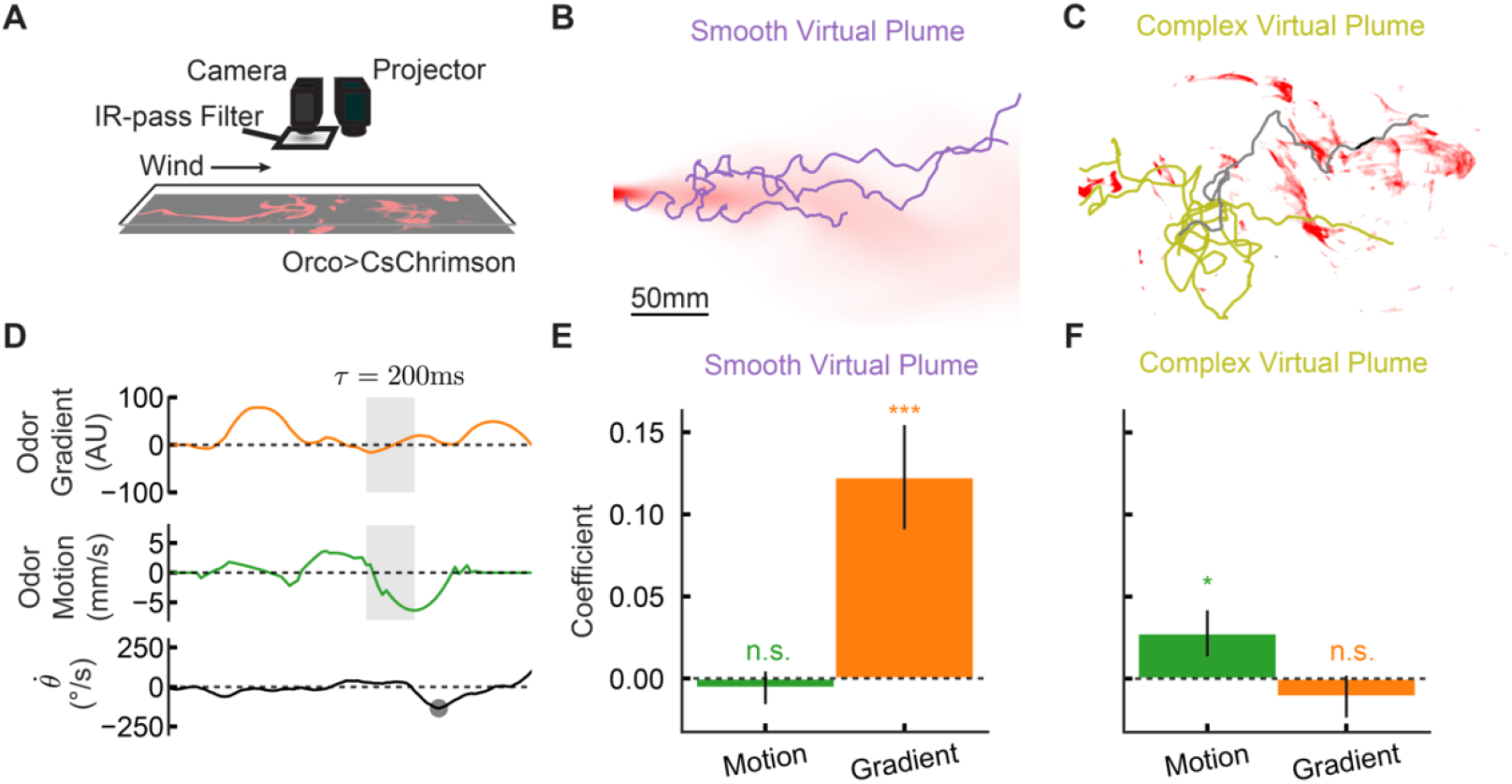
Flies make turns consistent with plume-appropriate directional odor cues. (A) Apparatus uses a projector to generate spatiotemporally patterned olfactory stimuli in an arena of walking flies. Olfactory stimuli were combined with 100mm/s laminar airflow across conditions. (B-C) Example trajectories from flies navigating the smooth (B) or complex (C) virtual plumes. (D) A time series from one highlighted trajectory (black path segment in panel (C)), plotting the fly’s angular velocity (black), odor gradient (orange), and odor motion (green) over several seconds. Saccade-like turns were detected as peaks in turn speed (gray dot). Gray boxes represent 200ms stimulus regions used to predict the direction of the upcoming turn. (E-F) Bar plots of logistic-regression coefficients (± cluster-robust SE) for flies navigating the smooth plume (E) or the complex plume (F). * indicates p < 0.05; *** indicates p < 0.005.

We tracked the trajectory and orientation of flies as they navigated the smooth and complex plume stimuli (**Fig. 5B-C**, see Methods). To estimate the gradient and motion signals that the flies experienced as they navigated the arena, we combined an estimate of each fly’s antennal location with a record of the projected stimulus (**Fig. 5D**). From each fly’s rotational velocities, we identified saccade-like turning events, which reoriented flies by ∼30° in the clockwise or counterclockwise direction, consistent with prior work (**Fig. 5D, Fig. S5**, see Methods) [33].

To isolate the relationships between olfactory directional cues and turns directed by the wind, we selected for further analysis time windows when flies were already facing upwind (± 20 degrees; see [46]). At these moments, rotations serve primarily to orient these flies in one crosswind direction or the other. We further restricted our analysis to flies that were engaged in upwind tracking in the bulk of the plume (see Methods). For turns that satisfied these criteria, we used stimulus features prior to turn onset to predict turn direction via logistic regression. In the smooth plume, the gradient cue but not the motion cue was a significant predictor (**Fig 5E**); conversely, in the complex plume the motion cue was a significant predictor, but the gradient cue was not **(Fig. 5F**). Thus, flies turn in response to different directional cues in different plumes, making crosswind turning decisions based on the most reliable cues in each plume type.

## Discussion

Odor gradient and odor motion sensing have been observed in multiple species, including both invertebrates [6, 7, 41-44, 46] and vertebrates [9, 13, 14]. We have shown that these cues provide complementary information about the plume centerline direction, with differential reliability in smooth and complex plumes. Specifically, gradients are reliable indicators of the centerline of smooth plumes and motion is a reliable indicator of the centerline of complex plumes.

Optimizing neural network models to determine centerline direction led these models to sense gradient and motion features in the smooth and complex plumes, respectively. When embedded in navigating agents, these feature-sensing models enhanced navigation performance only when navigators use them in the appropriate, plume-specific context. Fruit flies navigated virtual odor plumes in a manner that was consistent with these predictions: in smooth plumes, gradient cues predicted flies’ orienting decisions; in complex plumes, motion cues predicted flies’ orienting decisions.

Bilateral sensing can support navigation in different environments, in part because gradient and motion offer independent, complementary information to navigators. Recognizing this functional distinction may advance our understanding of bilateral odor processing principles in the brain. In the fruit fly, several neural pathways combine olfactory information across hemispheres, including olfactory receptor neurons themselves, which project bilaterally to the antennal lobes [43]. Recent experiments indicate that, downstream of the olfactory receptor neurons, projection neurons are sensitive to differences in odor intensity across the antennae [44]. Several lateral horn neurons also project bilaterally [54], and one such cell type has been implicated in gradient-driven behavior [53]. Although past research in these circuits has focused on gradient-coding [43, 44, 53], our work suggests that spatiotemporal coding principles in these and other bilaterally integrating olfactory circuits may support sensing multiple, distinct bilateral cues.

In this work, we tested behavioral predictions that we made in part by optimizing neural networks to solve navigational tasks using natural odor statistics. Goal-directed optimization of networks over natural stimuli is a powerful tool for understanding neural circuits [55]. In *Drosophila*, it has been used to explain functional properties of visual circuits [49, 56-58]. Here this approach provided a normative rationale for sensing both odor gradient and odor motion cues to resolve the direction to centerline across different plume conditions. This approach also provided trained filters that can be interpreted as a hypothesis about neural computation. In vision, similar optimization methods have produced neural networks with functional properties that predict experimentally measured neural responses [49, 58]. The filters trained in the complex plume exhibit a structure that is like the classical BL model of visual motion detection [50], combining immediate sensitivity to stimulation at one location with delayed, opposite-sign sensitivity to stimulation at a neighboring location (**Fig. 3**). Notably, differences in input statistics across plume types generated very different learned network architectures. Evidence in flies and other animals suggests that visual scene statistics strongly influence algorithms for visual motion detection [56, 59-64]. Exploring how the olfactory system is adapted processing odor signals in these diverse plume contexts may be an especially fruitful avenue for understanding complex neural computation.

Most broadly, our work indicates that directional odor cues can inform crosswind navigation across very different plumes. This insight motivates new avenues for theoretical and empirical research into odor navigation. Crosswind navigation appears more sophisticated and better informed by directional odor cues than previously realized. In diverse, naturalistic plumes, bilateral odor sensing provides more information than just the timing cues that gate upwind turns. Notably, bilateral odor sensing is widely conserved and supports navigation in invertebrates [6, 7, 41-44, 46, 65] and in vertebrates [9, 13, 14, 45, 47], including in humans [15]. Thus, bilateral and directional odor cues may be integral to odor plume navigation across animals. Moreover, our minimal network model is among the simplest that can extract gradient and motion information from bilateral timeseries of odor intensities. This model and others we studied are not fly-specific, beyond the spacing of the bilateral sensors. As a result, our computational approaches and results seem likely to be relevant to other species that use bilateral sensing for olfactory navigation.

## Acknowledgements

This work was funded by NIH R01 NS132840. GS was funded by a CAPES fellowship.

## Author contributions

The project was conceived by all authors. Numerical plume analysis and machine learning models were undertaken by BZ. Agent navigation and fly navigation measurements and analysis were performed by SB, with help from VJ and GS. The paper was written by SB, BZ, DAC, and TE.

---

**Figure S2:**
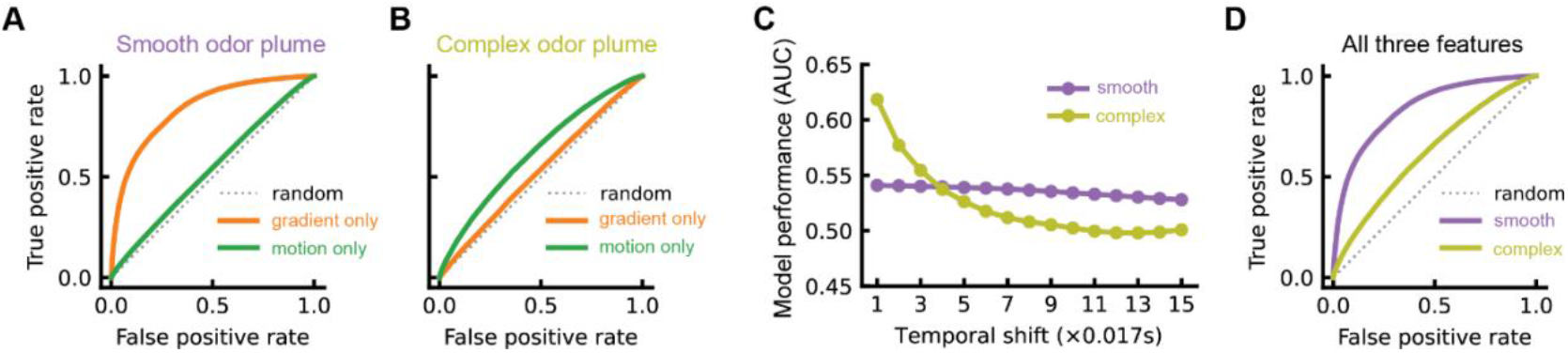
(A) Receiver operating curves associated with **Fig. 2B**. (B) Receiver operating curves associated with **Fig. 2C**. (C) ROC AUC scores as the functions of the temporal shift size in the engineered motion feature in both the smooth and complex plumes. (D) Receiver operating curves associated with **Fig. 2D, E**, with ROC AUC scores 0.847 (smooth plume) and 0.612 (complex plume).

**Figure S3:**
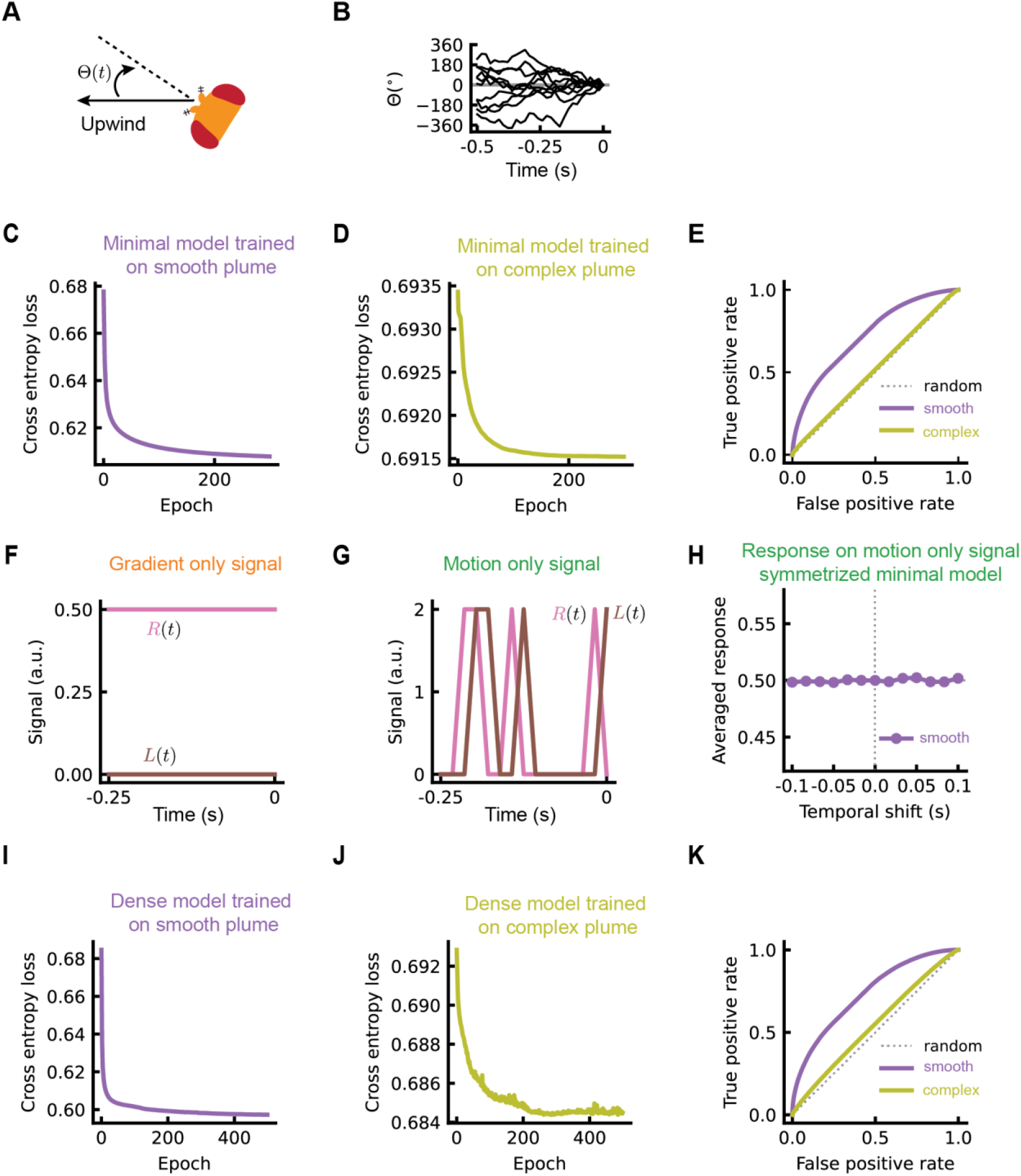
(A) The angle between the heading of the fly and the upwind direction is Θ(*t*). (B) Example trajectories of the angle Θ(*t*) in our simulations to generate synthetic traces for model training. At the time 0, when turning decision is made, the heading of the fly is always exactly upwind. (C, D) Training loss of the MNM as a function of the training epoch in the smooth (C) and complex (D) odor plumes. (E) Test data receiver operating curves for the MNMs trained in different plumes, with AUC scores 0.726 (smooth plume) and 0.517 (complex plume). (F) Example testing signals with only gradient information (related to **Fig. 3D, I**). (G) Example testing signals with only motion information, creating correlations between the antenna at a single temporal offset (related to **Fig. 3E, J**, see Methods). (H) Similar to Figure 3E, but with symmetrized filters for the MNM trained in the smooth plume. (I, J) Training loss of the DNM as a function of the training epoch in the smooth (I) and complex (J) odor plumes. (K) Test data receiver operating curves for the DNMs trained in different plume types with AUC scores 0.735 (smooth plume) and 0.543 (complex plume).

**Figure S4:**
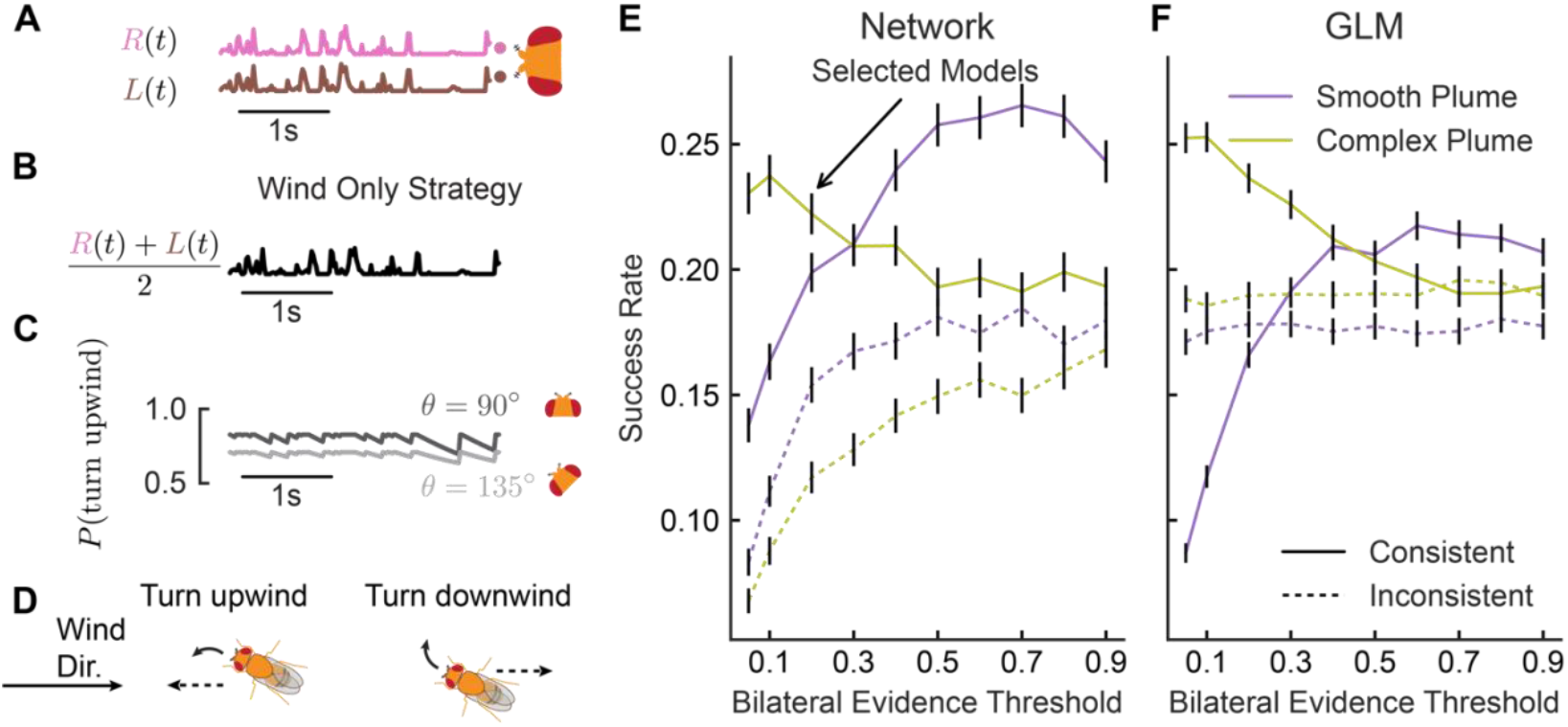
Outline of the baseline strategy and performances of different implementations of the bilateral strategy. (A) Example traces of bilateral signal intensity in a simulated agent. (B-D) In a baseline strategy that only uses wind as a directional cue, agents average the signal across the antennae (B). The agents calculate an upwind goal probability (C) based on this average intensity and on their current orientation.(D) After drawing an upwind or downwind goal direction on the basis of this computed probability, agents turned upwind or downwind according to the draw. This previously described model increases upwind turning in high intermittency and high frequency environments, and was parameterized on the basis of prior work [32]. (E-F) Agents that calculated directional odor features used the baseline model when the output of their directional odor model was weak. We tested performance across a range of thresholds on the strength of the bilateral evidence, both in agents that used neural network calculations(E) and in agents that used a GLM using predefined directional features (gradient and motion as in **Figure 2**) (F) as predictors of the centerline direction. Although these thresholds affect absolute performance levels, across thresholds agents using consistent features (the output of smooth-trained networks or smooth-trained, feature-based GLMs in the smooth plume; the output of complex-trained networks or complex-trained, feature-based GLMs in the complex plume) outperformed agents using inconsistent features (the output of complex-trained networks or complex trained GLMs in the smooth plume; the output of smooth-trained networks or smooth-trained GLMs in the complex plume).

**Figure S5:**
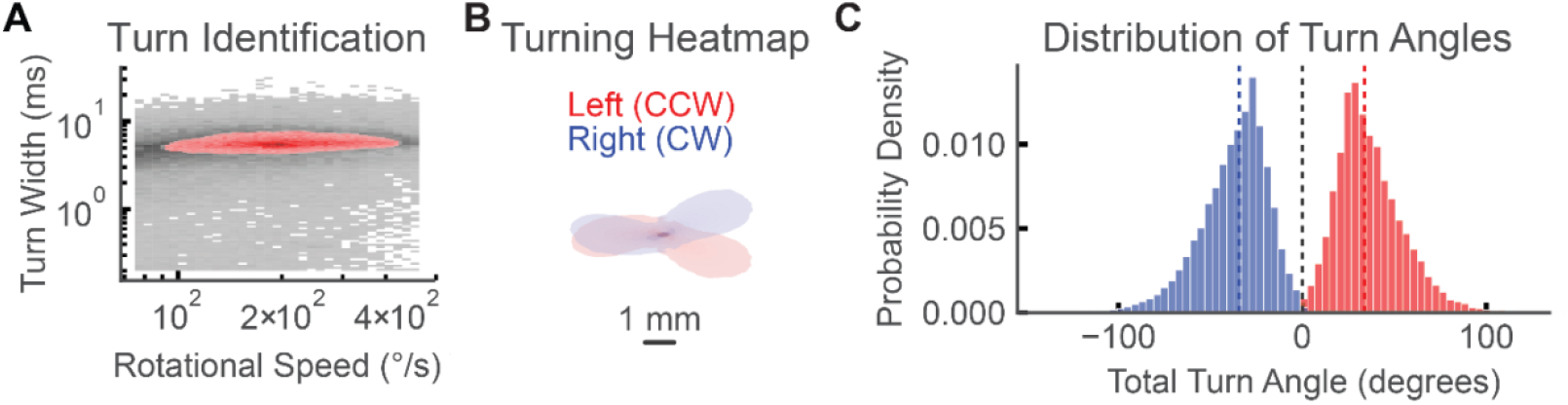
Detection and validation of saccade-like turns. (A) Histogram of the width (at half height) and amplitude of peaks in flies’ instantaneous rotational speed (log scale). An over-represented, dense region was isolated using a density based cluster-finding algorithm (DBSCAN, maroon) [66]. (B) Density of fly position in the vicinity (+/-200ms) of peaks labeled as turns. Positions are translated and rotated so they have identical spatial coordinates and orientations at the moment of the detected peak in rotational speed. Trajectories are colored according to the sign of the rotational velocity at the peak rotational speed. This density map represents the mean trajectories associated with left and right saccades. (C) Angular displacement during the identified turn events, colored according to the sign of the rotational velocity at the peak rotational speed. Median values (dashed lines; -34.2°, +33.8°) are similar to turn displacements reported in flies navigating plumes of smoke [33].

## Methods

### Plume movies

There were two types of plumes used in this paper, and we indicated them as smooth and complex plumes, respectively. The complex plume was a 60Hz movie with 3600 frames. We omitted the first 300 frames in order to make sure the odor plumes reached steady state and also omitted the last 1800 frames due to the gradual decaying of the intensity. Thus, only the rest 1500 frames were used. Each frame was a matrix of size 1088 × 1728, and each pixel was a 0.153 mm by 0.153 mm square. In the original video, the pixel values ranged from 0 to 120, and for consistency, we linearly rescaled them to the range from 0 to 255. The smooth plume was obtained from a published paper [20, 22], and we rescaled their original movie to match the time and space resolution of our sparse plume. Each frame was a matrix of size 1088 × 1696. The centerline of the smooth plume was set at 544 in vertical direction, while the centerline of the complex plume was set at 590 according to the exact location of the odor source.

For each movie, we calculated the averaged frame and used it to determine sampling regions and to avoid regions with averaged plume intensity below certain thresholds. After this, 1000 random locations were uniformly selected in the valid regions. For each location, we assumed there was an artificial fly with two antenna 3 pixels apart, and the signal into each antenna was filtered by a 2d Gaussian kernel with a zero mean and a standard deviation of 1.5 pixels. Each input to one antenna was a 30-timestep or 0.5s long vector. For each movie, we created two types of datasets. In the first dataset, the orientation of the fly was fixed, and the fly always faced upwind, which means that if the fly was in the lower half of the frame, the centerline was always to its right. In the second dataset, we allowed the orientation of the fly to undergo a Wiener process, but the data was collected in a way that in the last data point, the fly was facing upwind. Explicitly, the diffusion equation was

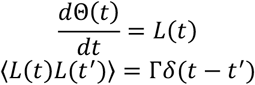

where the constant Γ was set to a value that the square root of the mean squared angular change in 0.5s was π. In this way, for each input vector, only the last few or most recent data points were informative about the centerline direction, since the fly could face any directions for data points that were further in the past.

### Logistic regression models

We engineered three distinct scalar features, and indicated them as odor sum, odor gradient, and odor motion, respectively. Suppose the filtered signals received by the left and right antenna were represented by *L*(*t*) and *R*(*t*), respectively, where *t* ranged from 0 to 29. The odor sum feature was obtained by the following equation:

The odor gradient feature:

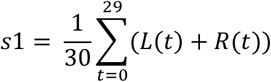

The odor motion feature:

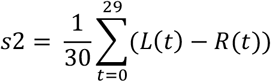

where τ varied from 1 to 15. In practice, τ = 1 gave us the best testing results. Each of the scalar features was fed into a Logistic regression model:

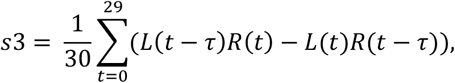

where σ(*x*) = 1/(1 + exp (−*x*)) was the sigmoid function and *b*_1_ was the intercept.

The dataset was generated with fly orientation fixed to face upwind. The training data contained 2,400,000 samples, and the testing data contained 600,000 samples. For each of the features, the logistic regression models were trained on 100 independently generated datasets to get enough statistics. The loss function was cross entropy.

### Artificial neural network model (ANN)

We built two types of ANNs: minimum ANN and dense ANN. In the minimum ANN model, there were two filters *f*_1_ and *f*_2_, and each was a vector with length 30. These 60 values and one bias term were the only trainable parameters in the model. The model was built as follows:

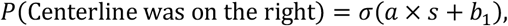

where *ReLU* was the rectified linear unit function and *b*_2_ was the intercept. The learning rate was set at 10^−4^, the batch size was 500 and the total number of training epochs was 500.

In the dense ANN, we used a 2-layer dense neural network with each layer 20 neurons. More complex dense models might result in overfitting. In this model, the input layer had a dimension of 60, concatenating the two input signals. Each neuron in the first hidden layer received a weighted sum of the input layer, and the weights could be seen as two temporal filters, each with a length of 30. Thus, there were 20 pairs of temporal filters in total as the first layer was mapped onto the first hidden layer. The learning rate was set at 10^−4^, the batch size was 500 and the total number of training epochs was 500.

The training data were generated with the fly orientation undergoing a Wiener process. For each type of datasets, there were 2,400,000 training samples and 600,000 testing samples. The loss function was the cross entropy.

### Testing trained models on synthetic stimuli

We designed two types of synthetic stimuli to test how the trained models respond to motion or gradient features. Each stimulus contained two signals, corresponding to the left and right antennae, respectively, and the time resolution was the same as in the training samples, which was 1/60 s. For the motion-feature stimuli, left signal had random binary inputs, 0 or 1, and the right signal was the shifted version of the left signal, and the shift size varied from -6 to 6. For the gradient-feature stimuli, both left and right signals were nonnegative constants, but their separation varied from -0.5 to 0.5 and the mean of the two signals was fixed at 0.25.

### Hassenstein Reichardt correlator

The simplest form of the Hassenstein Reichardt correlator can be written as

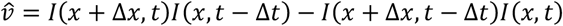

Expanding it to the second order gives

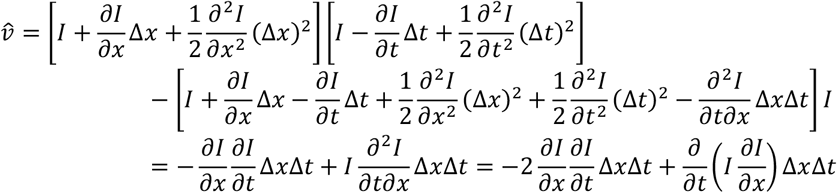

the first term of which (without the factor of 2) was used to calculate the motion signals in the Figure 1. The last term in the last line is a total derivative of time, or a surface term, and usually can be omitted when the velocity is estimated as an average over the time it takes for the filament to pass over the two antennae[67].

### Agent simulations

Agents navigated the plume movies described previously. They were initialized at random positions within a box downwind of the plume odor source. The box extended 200 to 250mm downwind of the source, and +/-83mm in the crosswind direction from the plume centerline. Initial agent orientation was also randomly initialized. At each movie timestep, agents moved forward at a speed of 10mm/s in the direction of their current orientation. Turns occurred stochastically, at times generated through a Poisson process with an average rate of 4/3 Hz. We imposed reflective spatial boundaries that kept agents inside the region of the plume movie (a ‘right’ wall at *x* = 270; a ‘top’ wall at *y* = 160; a ‘bottom’ wall at = 0). The upwind ‘left’ wall was not reflecting. Reaching *x* = 0 terminated trajectories. Agents navigated for 90s, or until their search was terminated by entering the goal region or passing upwind of the goal. For smooth plumes, the goal region was a 10mm circle centered at (0, 85). Because of a minor movie asymmetry, the goal region of the complex plume was centered at (0, 90).

When turns occurred, agent orientation changed by an absolute magnitude that was generated stochastically by sampling a normal distribution with a mean of 30° and standard deviation of 8°, following previous models of fly turning, and consistent with our measurements of behavior here (**Fig S5**) [33]. At each timestep, agents sensed stimulus intensity at two antennae spaced 306um apart along an axis perpendicular to agent orientation. In the *baseline strategy*, agents averaged these two intensity samples. This average intensity was combined with current orientation information into a goal vector according to a previously published model that describes flies’ upwind orientation decisions across a diverse array of timing statistics[32]. Specifically, this average was binarized based on whether it exceeded a noise floor (5AU). When the binarized signal was above threshold, an integrator signal, *R*, was set to 1. When the binarized signal fell below threshold, *R* decayed towards 0 with a 0.97s timescale. *R* was combined with agent orientation to generate an upwind turning bias:

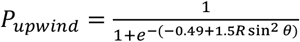

(parameters from [32]). Upwind turns were generated with probability *P*_*upwind*_ (and downwind turns occurred with the complementary probability 1 − *P*_*upwind*_).

In the *bilateral strategy*, sampled odor intensity values at the left and right antennae were scaled and preprocessed according to our neural network procedures (see prior section). 500ms histories of each signal were passed to our trained Dense Network Models to generate an estimated probability that the centerline was to the navigator’s left side 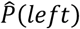. When 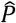 was near 0.5, agents using the bilateral strategy defaulted to the baseline strategy; otherwise, they turned left when 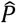 was large and turned right when 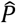 was small. Specifically, agents in Fig 4 used the baseline strategy when 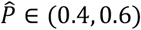.They turned right when 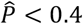 and turned right when 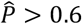.Performance at other thresholds is given is **Fig S4**. Baseline agents always used the baseline strategy.

### Fly strains and rearing

Fly rearing techniques were adapted from techniques in previous reports[32, 46]. We reared flies at 25ºC and 50% humidity under a 12 h–12 h light–dark cycle in plastic vials containing 10 ml standard glucose-cornmeal medium (Archon Scientific).

All flies in the study expressed a GMR-hid transgene that renders them blind. Optogenetic activation was achieved by expressing Chrimson (20X-UAS-CsChrimson) in Orco-expressing ORNs (Orco-GAL4). The genotypes used were: (1) w;gmr-hid;+ (gift from M. Murthy); (2) w;+;20XUAS-Chrimson (Bloomington, 55136); (3) w;+;Orco-Gal4 (gift from J. Carlson); (4) w;+.

### Behavioral apparatus

Flies navigated an optogenetic arena identical to the prior reports[32, 46], measuring 270mm (length) by 170mm (width) by 10mm (width). Laminar air (100mm/s) was introduced through an array of straws at the upwind side of the arena. A plastic mesh at the downwind side of the arena prevented flies from escaping.

Flies in the arena were illuminated with side-mounted 850 nm IR LED strips (Waveform Lighting) and recorded through an IR filter at 60 fps with a camera (FLIR Grasshopper USB 3.0). Dry air (Airgas) was passed through an array of aligned coffee straws to create 100mm/s laminar airflow through the arena. We used a projector (DLP LightCrafter 4500) to deliver red (627nm), csChrimson-activating light stimuli throughout the walking area. Stimuli updated at 60Hz.

The stimulus presentation and data recording operations were controlled in tandem by custom software written in Python 3.6.5. More detail about the control software is available in a prior publication [46].

### Stimulus design

We projected to-scale movies of the recorded smooth and complex plumes across the arena. Between the two plumes, data values do not reflect identical quantities (see Plume Data) and are not recorded on the same scales. We linearly scaled intensities in the smooth plume to span the range of intensities we could deliver through our projector. We used a transformation of the complex plume into projector intensities using the methods in [46]; this transformation includes a background subtraction to reduce noise in the movie, floor and ceiling thresholding of the signal, and then linear scaling to fill the projector dynamic range. The nonlinear thresholding process highlights contrast between odor filaments and interposed blank air.

Each trial lasted 2 minutes. On smooth-plume trials, we presented the first 2 minutes of the smooth movie; on complex-plume trials, the shorter complex movie was looped repeatedly to fill the 2-minute interval. A 1-minute rest period followed each trial. To avoid directional biases, each plume’s projection was flipped across its centerline between successive trials. Individual fly cohorts experienced either smooth-plume trials or complex-plume trials, but not both.

### Experimental Protocol

Between 10 and 30 females (<3 days post-eclosion) were placed in empty vials containing water-soaked cotton plugs at the bottom and top. These flies were starved for 3 days before recordings began. 24 hours before our experiments, the flies were fed 1 mM all trans-Retinal (ATR) (MilliporeSigma) dissolved in water and subsequently housed in the dark. All flies in a single starve vial were run in an experiment simultaneously.

We used the incubator light/dark cycle to identify “dawn” epochs (the first 3 hours of the light cycle) and “dusk” epochs (the last 3 hours of the light cycle). We ran all experiments inside these circadian windows, when fly activity peaks.

At the start of each experiment a fly cohort was aspirated into the back of the arena through a sealable hole in the arena’s ceiling. Flies acclimatized to the environment, including the laminar airflow, for 2 minutes prior to experiment onset. Each fly cohort navigated either smooth plume trials or complex plume trials. An experiment on one fly cohort consisted of 8 navigation trials.

### Preprocessing

We identified fly head and abdomen positions from recorded videos using SLEAP [68]. We smoothed position trajectories using a 4^th^ order Savitsky-Golay filter with a 21-sample window (0.3s). The analytic derivatives of these polynomial fits were used to quantify x and y velocities. To determine fly orientation, we first estimated its body outline using scikit-image’s “canny” function. We found the best-fit ellipse approximation to this outline and treated the major axis as the head-to-tail axis. When the fly was moving, our estimates of the ground velocity were used to disambiguate the head from the tail, since flies typically move forward rather than backward.

When the flies were still, we used the major axis orientation that best matched the abdomen-to-head angle found through SLEAP. We used the same smoothing operations on orientation traces to obtain smoothed fly orientation and rotational velocity traces.

To estimate the stimuli flies experienced, we adapted previously published gradient and motion calculation methods [46]. We estimated the antennal sensory axis as perpendicular to the major fly orientation axis and located it 1.5mm anterior to the fly abdomen location. Following a prior procedure [46], we accessed 15 pixels along the antenna axis of the fly at each time point and used the slope of projector intensity values across this slice as our gradient measure. Odor motion was estimated by cross-correlating slices from consecutive frames, identifying the lateral offset that maximized the correlation, and converting that offset to a velocity-like measure based on calibration (mm per pixel) and sampling frequency. These signals were then smoothed using an exponential filter (τ = 200ms) and z-scored for subsequent analysis.

### Behavioral Analysis

We located candidate turning events by finding local maxima in absolute angular velocity. A density-based clustering algorithm (DBSCAN algorithm; eps=0.1, min_samples=900) was applied to the joint distribution of log-transformed peak amplitude and peak height. We treated a dense region in the resulting distribution as exemplary saccade-like turns after inspecting the surrounding trajectories and the associated heading changes (**Fig. S5**).

For analysis, we retained only turns that met the following conditions:

1. Upwind orientation: Heading angles within ±20° of directly upwind (160–200°).
2. Active upwind walking: Positive horizontal velocity as determined via a fourth-order Butterworth filter (cutoff=0.2 Hz) applied to *vx*.
3. Arena region: Horizontal positions 5cm<*x*<25cm, with vertical position *y* contained within a 4cm band around the plume center line.

We modeled turn direction (left vs. right) using logistic regression. We used the smoothed and z-scored estimates of odor intensity, gradient, and motion as predictors. Specifically, we used their values 50ms prior to turn onset (time of crossing 20% turn speed peak height on the rising edge of the peak). 50ms was chosen to reflect an approximate minimal response time to bilateral odor cues [43]. Separate models were fit for each plume type. Observations from each fly trajectory were grouped to compute cluster-robust (trajectory-level) standard errors in the Statsmodels logistic framework. This approach provided estimates of how each odor-derived cue influenced the likelihood of turning left or right under controlled upwind conditions.

## Notes

### Competing Interest Statement

The authors have declared no competing interest.

## References

1. Flugge, C., Geruchliche raumorientierung von Drosophila melanogaster. Zeitschrift f \ u r vergleichende Physiologie, 1934. 20(4): p. 463–-500.

2. Murlis, J.E. JS; Carde, RT, Odor plumes and how insects use them. Annual review of entomology, 1992. 37(1): p. 505–532.

3. Mafra-Neto, A. and R. Cardé, Influence of plume structure and pheromone concentration on upwind flight of Cadra cautella males. Physiological Entomology, 1995. 20(2): p. 117–133.

4. Mafra-Neto, A. and R.T. Cardé, Fine-scale structure of pheromone plumes modulates upwind orientation of flying moths. Nature, 1994. 369(6476): p. 142–144.

5. Lockery, S.R., The computational worm: spatial orientation and its neuronal basis in C. elegans. Curr Opin Neurobiol, 2011. 21(5): p. 782–90.

6. Reeder, P.B. and B.W. Ache, Chemotaxis in the Florida spiny lobster, Panulirus argus. Animal Behaviour, 1980. 28(3): p. 831–839.

7. Atema, J., Eddy Chemotaxis and Odor Landscapes: Exploration of Nature With Animal Sensors. Biol Bull, 1996. 191(1): p. 129–138.

8. Webster, D. and M. Weissburg, Chemosensory guidance cues in a turbulent chemical odor plume. Limnology and Oceanography, 2001. 46(5): p. 1034–1047.

9. Gardiner, J.M. and J. Atema, The function of bilateral odor arrival time differences in olfactory orientation of sharks. Curr Biol, 2010. 20(13): p. 1187–91.

10. Scholz, A.T., et al., Imprinting to chemical cues: the basis for home stream selection in salmon. Science, 1976. 192(4245): p. 1247–1249.

11. Nevitt, G.A., Olfactory foraging by Antarctic procellariiform seabirds: life at high Reynolds numbers. Biol Bull, 2000. 198(2): p. 245–53.

12. Wallraff, H.G., Avian olfactory navigation: its empirical foundation and conceptual state. Animal Behaviour, 2004. 67(2): p. 189–204.

13. Khan, A.G., M. Sarangi, and U.S. Bhalla, Rats track odour trails accurately using a multi-layered strategy with near-optimal sampling. Nature communications, 2012. 3(1): p. 703.

14. Catania, K.C., Stereo and serial sniffing guide navigation to an odour source in a mammal. Nat Commun, 2013. 4: p. 1441.

15. Porter, J., et al., Mechanisms of scent-tracking in humans. Nat Neurosci, 2007. 10(1): p. 27–9.

16. Martin, J.P., et al., The neurobiology of insect olfaction: sensory processing in a comparative context. Progress in neurobiology, 2011. 95(3): p. 427–447.

17. Baker, K.L., et al., Algorithms for Olfactory Search across Species. J Neurosci, 2018. 38(44): p. 9383–9389.

18. Szyszka, P., T. Emonet, and T.L. Edwards, Extracting spatial information from temporal odor patterns: Insights from insects. Current Opinion in Insect Science, 2023. 59: p. 101082.

19. Emonet, T. and M. Vergassola, Olfactory cues and memories in animal navigation. Nature Reviews Physics, 2024. 6(4): p. 215–216.

20. Connor, E.G., M.K. McHugh, and J.P. Crimaldi, Quantification of airborne odor plumes using planar laser-induced fluorescence. Experiments in Fluids, 2018. 59: p. 1–11.

21. Reddy, G., V.N. Murthy, and M. Vergassola, Olfactory sensing and navigation in turbulent environments. Annual Review of Condensed Matter Physics, 2022. 13(1): p. 191–213.

22. Alvarez-Salvado, E., et al., Elementary sensory-motor transformations underlying olfactory navigation in walking fruit-flies. Elife, 2018. 7.

23. Keller, T.A. and M.J. Weissburg, Effects of odor flux and pulse rate on chemosensory tracking in turbulent odor plumes by the blue crab, Callinectes sapidus. The Biological Bulletin, 2004. 207(1): p. 44–55.

24. Geisler, W.S., Visual perception and the statistical properties of natural scenes. Annu. Rev. Psychol., 2008. 59(1): p. 167–192.

25. Celani, A., E. Villermaux, and M. Vergassola, Odor landscapes in turbulent environments. Physical Review X, 2014. 4(4): p. 041015.

26. Riffell, J.A., L. Abrell, and J.G. Hildebrand, Physical processes and real-time chemical measurement of the insect olfactory environment. J Chem Ecol, 2008. 34(7): p. 837–53.

27. Taylor, G.I., Diffusion by continuous movements. Proceedings of the london mathematical society, 1922. 2(1): p. 196–212.

28. Simoncelli, E.P. and B.A. Olshausen, Natural image statistics and neural representation. Annual review of neuroscience, 2001. 24(1): p. 1193–1216.

29. Vickers, N.J., Mechanisms of animal navigation in odor plumes. Biol Bull, 2000. 198(2): p. 203–12.

30. Carde, R.T. and M.A. Willis, Navigational strategies used by insects to find distant, windborne sources of odor. J Chem Ecol, 2008. 34(7): p. 854–66.

31. Carde, R.T., Navigation Along Windborne Plumes of Pheromone and Resource-Linked Odors. Annu Rev Entomol, 2021. 66: p. 317–336.

32. Jayaram, V., et al., Temporal novelty detection and multiple timescale integration drive Drosophila orientation dynamics in temporally diverse olfactory environments. PLoS Comput Biol, 2023. 19(5): p. e1010606.

33. Demir, M., et al., Walking Drosophila navigate complex plumes using stochastic decisions biased by the timing of odor encounters. Elife, 2020. 9.

34. van Breugel, F. and M.H. Dickinson, Plume-tracking behavior of flying Drosophila emerges from a set of distinct sensory-motor reflexes. Curr Biol, 2014. 24(3): p. 274–86.

35. Budick, S.A. and M.H. Dickinson, Free-flight responses of Drosophila melanogaster to attractive odors. J Exp Biol, 2006. 209(Pt 15): p. 3001–17.

36. Jayaram, V., N. Kadakia, and T. Emonet, Sensing complementary temporal features of odor signals enhances navigation of diverse turbulent plumes. Elife, 2022. 11.

37. Rigolli, N., et al., Alternation emerges as a multi-modal strategy for turbulent odor navigation. Elife, 2022. 11.

38. Verano, K.V.B., E. Panizon, and A. Celani, Olfactory search with finite-state controllers. Proceedings of the National Academy of Sciences, 2023. 120(34): p. e2304230120.

39. Ouyang, B., et al., Simple olfactory navigation in air and water. J Theor Biol, 2024. 595: p. 111941.

40. Balkovsky, E. and B.I. Shraiman, Olfactory search at high Reynolds number. Proc Natl Acad Sci U S A, 2002. 99(20): p. 12589–93.

41. Borst, A. and M. Heisenberg, Osmotropotaxis in Drosophila melanogaster. Journal of comparative physiology, 1982. 147(4): p. 479–484.

42. Duistermars, B.J., D.M. Chow, and M.A. Frye, Flies require bilateral sensory input to track odor gradients in flight. Current Biology, 2009. 19(15): p. 1301–1307.

43. Gaudry, Q., et al., Asymmetric neurotransmitter release enables rapid odour lateralization in Drosophila. Nature, 2013. 493(7432): p. 424–8.

44. Taisz, I., et al., Generating parallel representations of position and identity in the olfactory system. Cell, 2023. 186(12): p. 2556–2573. e22.

45. Liu, A., et al., Mouse Navigation Strategies for Odor Source Localization. Front Neurosci, 2020. 14: p. 218.

46. Kadakia, N., et al., Odour motion sensing enhances navigation of complex plumes. Nature, 2022. 611(7937): p. 754–761.

47. Rajan, R., J.P. Clement, and U.S. Bhalla, Rats smell in stereo. Science, 2006. 311(5761): p. 666–70.

48. Fitzgerald, J.E., et al., Symmetries in stimulus statistics shape the form of visual motion estimators. Proceedings of the National Academy of Sciences, 2011. 108(31): p. 12909–12914.

49. Mano, O., et al., Predicting individual neuron responses with anatomically constrained task optimization. Curr Biol, 2021. 31(18): p. 4062–4075 e4.

50. Barlow, H.B. and W.R. Levick, The mechanism of directionally selective units in rabbit’s retina. J Physiol, 1965. 178(3): p. 477–504.

51. Salazar-Gatzimas, E., et al., Direct Measurement of Correlation Responses in Drosophila Elementary Motion Detectors Reveals Fast Timescale Tuning. Neuron, 2016. 92(1): p. 227–239.

52. Rigolli, N., et al., Learning to predict target location with turbulent odor plumes. Elife, 2022. 11: p. e72196.

53. Mohamed, A.A.M., B.S. Hansson, and S. Sachse, Third-Order Neurons in the Lateral Horn Enhance Bilateral Contrast of Odor Inputs Through Contralateral Inhibition in Drosophila. Front Physiol, 2019. 10: p. 851.

54. Dolan, M.J., et al., Neurogenetic dissection of the Drosophila lateral horn reveals major outputs, diverse behavioural functions, and interactions with the mushroom body. Elife, 2019. 8.

55. Turner, M.H., et al., Stimulus-and goal-oriented frameworks for understanding natural vision. Nature neuroscience, 2019. 22(1): p. 15–24.

56. Wu, N., et al., Broken time-reversal symmetry in visual motion detection. Proceedings of the National Academy of Sciences, 2025. 122(10): p. e2410768122.

57. Zhou, B., et al., Shallow neural networks trained to detect collisions recover features of visual loom-selective neurons. Elife, 2022. 11: p. e72067.

58. Lappalainen, J.K., et al., Connectome-constrained networks predict neural activity across the fly visual system. Nature, 2024. 634(8036): p. 1132–1140.

59. Chen, J., et al., Asymmetric ON-OFF processing of visual motion cancels variability induced by the structure of natural scenes. Elife, 2019. 8: p. e47579.

60. Clark, D.A. and J.E. Fitzgerald, Optimization in visual motion estimation. Annual Review of Vision Science, 2024. 10.

61. Clark, D.A., et al., Flies and humans share a motion estimation strategy that exploits natural scene statistics. Nature neuroscience, 2014. 17(2): p. 296–303.

62. Fitzgerald, J.E. and D.A. Clark, Nonlinear circuits for naturalistic visual motion estimation. Elife, 2015. 4: p. e09123.

63. Leonhardt, A., et al., Asymmetry of Drosophila ON and OFF motion detectors enhances real-world velocity estimation. Nature neuroscience, 2016. 19(5): p. 706–715.

64. Yildizoglu, T., et al., A neural representation of naturalistic motion-guided behavior in the zebrafish brain. Current Biology, 2020. 30(12): p. 2321–2333. e6.

65. Louis, M., et al., Bilateral olfactory sensory input enhances chemotaxis behavior. Nature neuroscience, 2008. 11(2): p. 187–199.

66. Ester, M., et al. A density-based algorithm for discovering clusters in large spatial databases with noise. in kdd. 1996.

67. Sinha, S.R., W. Bialek, and R.R.D.R. Van Steveninck, Optimal local estimates of visual motion in a natural environment. Physical review letters, 2021. 126(1): p. 018101.

68. Pereira, T.D., et al., SLEAP: A deep learning system for multi-animal pose tracking. Nature methods, 2022. 19(4): p. 486–495.

